## Supplementary Note for "Sharing information between related diseases using Bayesian joint fine mapping increases accuracy and identifies novel associations in six immune mediated diseases"

February 13, 2019

#### 1 LD structures corresponding to joint tagging

We consider 3 SNPs. SNPs 1 and 2 are causal and SNP 3 is not causal. The LD correlation matrix between the SNPs is

$$\Sigma = \begin{bmatrix} 1 & r_{12} & r1 \\ r_{12} & 1 & r2 \\ r1 & r2 & 1 \end{bmatrix}$$

Because $\Sigma$ is a correlation matrix, it must be positive definite, which means $r_{12}, r_1, r_2$ must satisfy

$$-2 * r_1 * r_2 * r_{12} + r_1^2 + r_2^2 + r_{12}^2 \leq 1 \qquad (1)$$

so that $r_{12}$, $r_1, r_2$ are constrained to lie within an ellipse. If the true expected $Z$ scores from a joint model against all SNPs is $Z_J = (\zeta_1, \zeta_2, 0)'$, then the expected marginal $Z$ scores are $Z_M \simeq \Sigma Z_J = (z_1, z_2, z_3)'$ where $z_i = E\left(\frac{\beta_i}{\sigma_i}\right)$, and $\beta_i, \sigma_i$ are the log odds ratio and its standard error, respectively, for SNP $i$. The region within the ellipse that correspond to joint tagging is defined by the intersection of

$$z_3 \simeq |\zeta_1 r_1 + \zeta_2 r_2| > |\zeta_1 + \zeta_2 r_{12}| \simeq z_1$$

$$z_3 \simeq |\zeta_1 r_1 + \zeta_2 r_2| > |\zeta_2 + \zeta_1 r_{12}| \simeq z_2$$

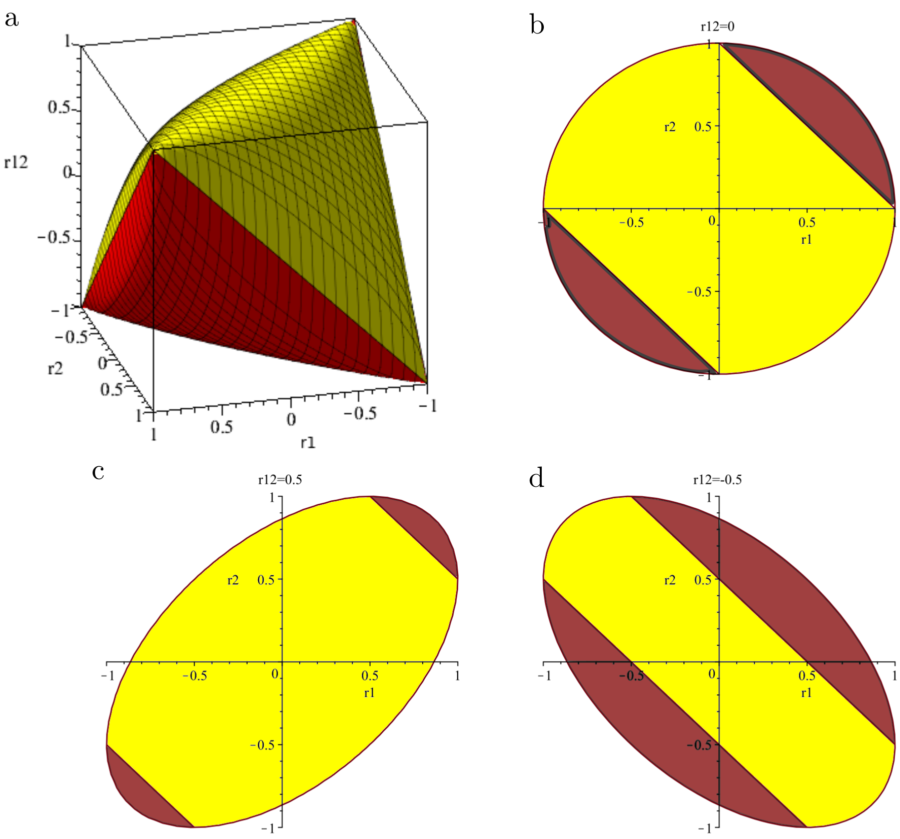


Figure 1: The 3-way correlations between 2 causal SNPs ($r_{12}$, z axis) and each causal SNP and a potential joint tag ($r_1$, and $r_2$, x and y axes respectively) lie in a simplex. **a** Assuming $\zeta_1 = \zeta_2$, then points within this simplex may be coloured according to whether joint tagging is expected (red) or not (yellow) if both causal variants have equal effect sizes. **b**, **c**, **d** show planes through this simplex when the causal variants are uncorrelated ($r_{12} = 0$), positively correlated ($r_{12} = 0.5$) and negatively correlated ($r_{12} = -0.5$) respectively.

There are then 5 unknown parameters which control whether tagging is expected. However, we can make inference under some simplified expectations. For example, if we assume that the two causal SNPs have equal effect sizes measured by Z scores, ie $\zeta_1 = \zeta_2 \Rightarrow z_1 = z_2$, then this reduces to

$$|r_1 + r_2| > |1 + r_{12}|. \tag{2}$$

Equation (2) thus defines the subset of points within the simplex (within which these 3-way correlations must lie) which correspond to joint tagging under the assumption $\zeta_1 = \zeta_2$. In particular, note that this subset is non-empty - ie joint tagging is possible - even for unlinked causal variants, since setting $r_{12} = 0$ joint tagging requires a solution to the simultaneous inequalities derived from (1), (2):

$$\begin{aligned} |r_1 + r_2| &> 1 \\ r_1^2 + r_2^2 &\leq 1 \end{aligned} \tag{3}$$

and many such solutions exist, e.g. $r_1 = r_2 > 0.5$.

Alternatively, it may be more realistic to assume that the SNPs have equal odds ratios, $\beta^*$. Noting that $\sigma_i^2 \propto f_i(1 - f_i)$ where $f_i$ is the MAF of SNP $i$, we have joint tagging when

$$\begin{aligned} z_3 \simeq \left| \frac{\beta^*}{\sqrt{f_1(1-f_1)}} r_1 + \frac{\beta^*}{\sqrt{f_2(1-f_2)}} r_2 \right| &> \left| \frac{\beta^*}{\sqrt{f_1(1-f_1)}} + \frac{\beta^*}{\sqrt{f_2(1-f_2)}} r_{12} \right| \simeq z_1 \\ z_3 \simeq \left| \frac{\beta^*}{\sqrt{f_1(1-f_1)}} r_1 + \frac{\beta^*}{\sqrt{f_2(1-f_2)}} r_2 \right| &> \left| \frac{\beta^*}{\sqrt{f_1(1-f_1)}} r_{12} + \frac{\beta^*}{\sqrt{f_2(1-f_2)}} \right| \simeq z_2 \end{aligned}$$

which reduces to

$$\begin{aligned} z_3 \simeq \left| \frac{r_1}{\sqrt{f_1(1-f_1)}} + \frac{r_2}{\sqrt{f_2(1-f_2)}} \right| &> \left| \frac{1}{\sqrt{f_1(1-f_1)}} + \frac{r_{12}}{\sqrt{f_2(1-f_2)}} \right| \simeq z_1 \\ z_3 \simeq \left| \frac{r_1}{\sqrt{f_1(1-f_1)}} + \frac{r_2}{\sqrt{f_2(1-f_2)}} \right| &> \left| \frac{r_{12}}{\sqrt{f_1(1-f_1)}} + \frac{1}{\sqrt{f_2(1-f_2)}} \right| \simeq z_2 \end{aligned} \tag{4}$$

Decisions on whether individual observations corresponded to joint tagging in Fig. 2b–c were made on the basis of equations (3)-(4).

#### 2 Statistical inference of joint versus tag models

In this section, we consider how statistical inference will perform, when comparing joint models to tag models, by evaluating their likelihood ratio and the Bayesian Information Criterion (BIC). Let

$Z^o$ be the observed $Z$ scores for the joint 3-SNP model, with $Z^o \sim N(Z_M, \Sigma)$, $Z_M = \Sigma Z_J$.

$$H_0 : Z_J = \tilde{z}_J = (0, 0, \tilde{\zeta})'$$
$$H_1 : Z_J = z_j = (\zeta_1, \zeta_2, 0)'$$

For simplicity, we assume the MLE of the effect sizes under the appropriate hypothesis are their true values, i.e. $\hat{\zeta}_i = \zeta_i$, $i = 1, 2$, assuming $H_1$, and $\hat{\tilde{\zeta}} = r_1\zeta_1 + r_2\zeta_2$, assuming $H_0$. We are interested in evaluating the plausibility of situations in which we would erroneously infer $H_0$ when $H_1$ is true. Assuming a likelihood ratio is used for comparison, this would correspond to

$$\frac{\ell(Z_J = (\zeta_1, \zeta_2, 0)'|z_o)}{\ell(Z_J = (0, 0, \tilde{\zeta})'|z_o)} < 1,$$

where the left-hand side is the likelihood ratio of observing $Z$ scores $(z_1^o, z_2^o, z_3^o)$ under $H_1$ compared to $H_0$. Since $Z^o$ is Normally distributed, the inequality becomes

$$\frac{(2\pi|\Sigma|)^{-1/2} \exp\left(-\frac{1}{2}(z_o - z_m)'\Sigma^{-1}(z_o - z_m)\right)}{(2\pi|\Sigma|)^{-1/2} \exp\left(-\frac{1}{2}(z_o - \tilde{z}_m)'\Sigma^{-1}(z_o - \tilde{z}_m)\right)} < 1,$$

where $z_m = \Sigma z_J$ and $\tilde{z}_m = \Sigma \tilde{z}_J$. Simplifying, we obtain

$$z_m'\Sigma^{-1}z_m - \tilde{z}_m'\Sigma^{-1}\tilde{z}_m - 2z_o'\Sigma^{-1}(z_m - \tilde{z}_m) > 0$$
$$\Rightarrow \qquad z_J'\Sigma z_J - \tilde{z}_J'\Sigma\tilde{z}_J - 2z_o'(z_J - \tilde{z}_J) > 0,$$

where we used the fact that $\Sigma$ is symmetric and expressions for $z_m$ and $\tilde{z}_m$. Substituting $z_J$, $\tilde{z}_J$ and $\Sigma$ we obtain

$$2(\tilde{\zeta}z_3^o - \zeta_1 z_1^o - \zeta_2 z_2^o) + \zeta_1^2 + \zeta_2^2 - \tilde{\zeta}^2 + 2\zeta_1\zeta_2 r_{12} > 0. \tag{5}$$

We need to determine whether this condition can be satisfied, and if so, what is the probability of it being satisfied if $H_1$ is true.

The relative plausibility of the two models can also be assessed using Bayesian information criterion (BIC). Recall that the BIC of a model with $k$ parameters and based on $n$ sample points is $k\ln(n) - 2\hat{\ell}$, where $\hat{\ell}$ is the maximized log-likelihood of the model. Hence, when using BIC, in order for the model under which SNP 3 is causal to be preferred to the model under which SNPs 1 and 2 are causal, the following has to hold

$$\ln(n) - 2\hat{\ell}(Z_J = (0, 0, \tilde{\zeta})' | z_o) < 2\ln(n) - 2\hat{\ell}(Z_J = (\zeta_1, \zeta_2, 0)' | z_o).$$

This gives the following condition

$$2(\tilde{\zeta} z_3^o - \zeta_1 z_1^o - \zeta_2 z_2^o) + \zeta_1^2 + \zeta_2^2 - \tilde{\zeta}^2 + 2\zeta_1\zeta_2 r_{12} > -\ln(n). \tag{6}$$

We now proceed to evaluate probabilities of conditions (5) and (6). We assume $\zeta_1 = \zeta_2 := \zeta$ and $\tilde{\zeta} = \zeta(r_1 + r_2)$. Hence, $Z_J = (\zeta, \zeta, 0)'$ and we have

$$Z^o \sim N\left(\begin{bmatrix} \zeta(1+r_{12}) \\ \zeta(1+r_{12}) \\ \zeta(r_1+r_2) \end{bmatrix}, \Sigma\right),$$

Set

$$W := \tilde{\zeta} z_3^o - \zeta_1 z_1^o - \zeta_2 z_2^o$$

and note that $W$ is Normally distributed and (5) becomes

$$2W - \zeta^2(r_1 + r_2)^2 + 2\zeta^2(1 + r_{12}) > 0, \tag{7}$$

with

$$\mathbb{E}(W) = \zeta^2(r_1 + r_2)^2 - 2\zeta^2(1 + r_{12}) := -\sigma_W^2$$
$$\mathrm{Var}(W) = -\zeta^2(r_1 + r_2)^2 + 2\zeta^2(1 + r_{12}) = \sigma_W^2.$$

Condition $\mathrm{Var}(W) > 0$ evaluates to

$$r_1 + r_2 < \sqrt{2(1 + r_{12})}. \tag{8}$$

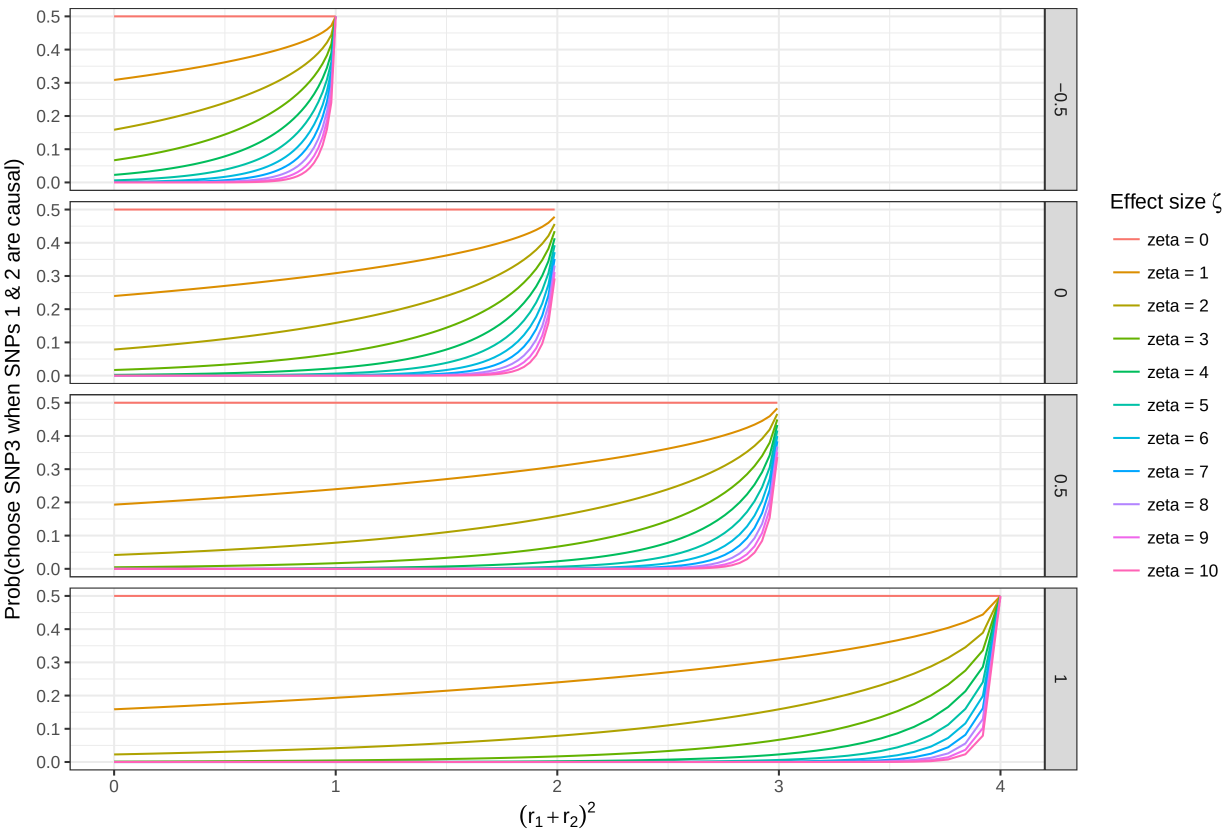


Figure 2: Probability of choosing a model under which SNP 3 is causal, when SNPs 1 and 2 are actually causal, $P(\zeta, r_1, r_2, r_{12})$, under likelihood ratio setup for varying values of $(r_1+r_2)^2$, squared sum of correlations of SNPs 1 and 2 with SNP 3, different effect sizes $\zeta$, and correlation $r_{12}$ between SNPs 1 and 2 (side panels). Note that there are no combinations of $r_1$ and $r_2$ satisfying conditions (1) and (8) when $r_{12} = -1$. $P(\zeta, r_1, r_2, r_{12})$ is bounded above by 0.5 and remains very small for $\zeta > 3$ until $(r_1 + r_2)^2$ is close to its maximum value.

We can now evaluate the probability of event (7), $P(\zeta, r_1, r_2, r_{12})$, as

$$P(\zeta, r_1, r_2, r_{12}) = \mathbb{P}(2W + \sigma_W^2 > 0) = 1 - \Phi(1/2\sigma_W)$$
$$= 1 - \Phi\left(\frac{1}{2}|\zeta|\sqrt{2(1 + r_{12}) - (r_1 + r_2)^2}\right).$$

It follows that the probability of choosing $H_0$ when $H_1$ is in fact true, is always less than 0.5. Additionally this probability decreases with the absolute value of effect size $\zeta$ and correlation between SNPs 1 and 2, $r_{12}$, and increases with the squared sum of correlations of causal SNPs 1 and 2 with the tagging SNP 3 (see Figure 2).

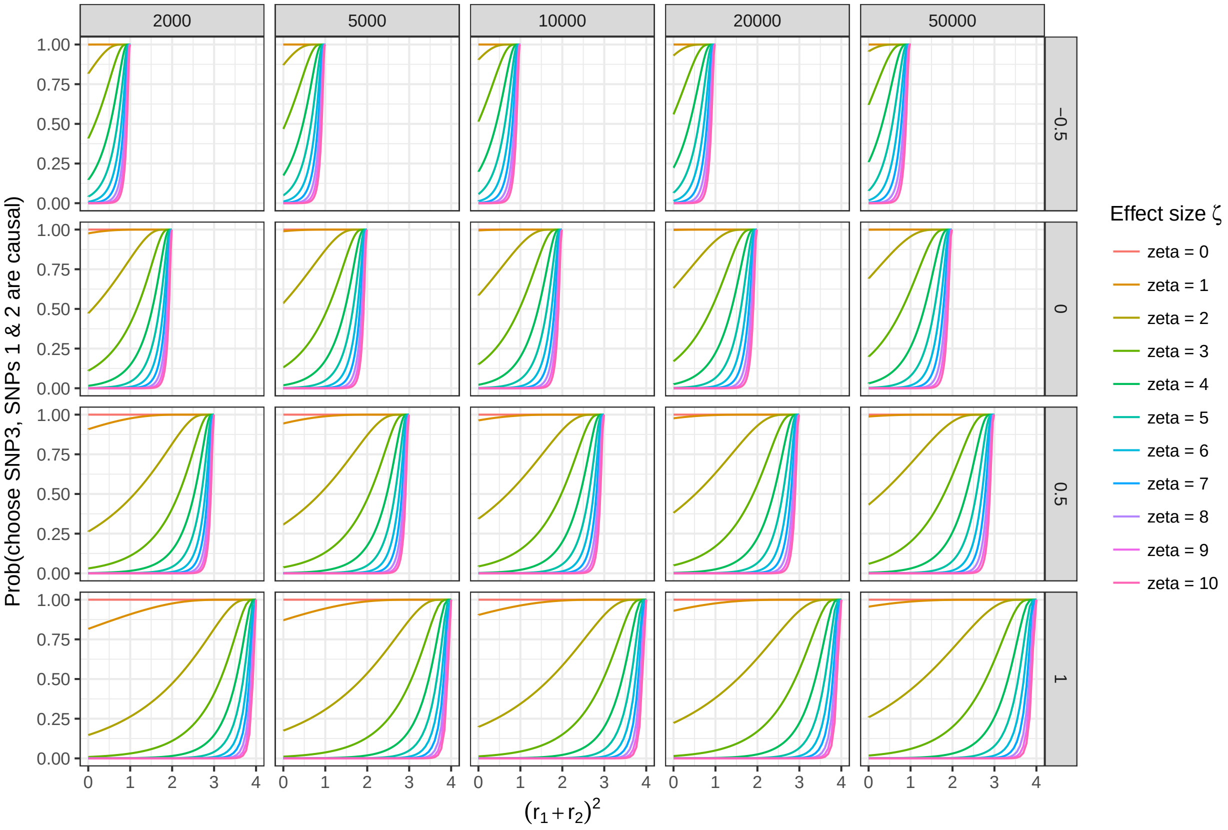


Figure 3: Probability of choosing a model under which SNP 3 is causal, when SNPs 1 and 2 are actually causal, $P(\zeta, r_1, r_2, r_{12}, n)$, under BIC setup for varying values of $(r_1 + r_2)^2$, squared sum of correlations of SNPs 1 and 2 with SNP 3, different effect sizes $\zeta$, correlation $r_{12}$ between SNPs 1 and 2 (side panels), and sample size $n$ (top panels). Note that there are no combinations of $r_1$ and $r_2$ satisfying conditions (1) and (8) when $r_{12} = -1$. $P(\zeta, r_1, r_2, r_{12}, n)$ remains very small for $\zeta > 3$ until $(r_1 + r_2)^2$ is close to its maximum value.

Similarly, the probability of the BIC condition (6), $P(\zeta, r_1, r_2, r_{12}, n)$, can be calculated to be

$$P(\zeta, r_1, r_2, r_{12}, n) = \mathbb{P}(2W + \sigma_W^2 > -\ln n) = 1 - \Phi\left(\frac{1}{2}\left(\sigma_W - \frac{1}{\sigma_M}\ln(n)\right)\right).$$

Again, the probability of choosing a SNP 3 model, when SNPs 1 and 2 are causal, is non-zero but is no longer bounded above. The general trends remain the same—$P(\zeta, r_1, r_2, r_{12}, n)$ increases with $(r_1+r_2)^2$ and decreases with $\zeta$ and $r_{12}$ (although the pool of admissible values $(r_1, r_2)$ increases with $r_{12}$). Additionally $P(\zeta, r_1, r_2, r_{12}, n)$ slightly increases with the sample size $n$ (see Figure 3).

#### 2.1 SNP 3 is causal

If $H_0$ were in fact true, we would be much less likely to erroneously infer $H_1$, as shown below. Assume $\zeta_1 = \tilde{\zeta} r_1$ and $\zeta_2 = \tilde{\zeta} r_2$. We have

$$Z^o \sim N\left(\begin{bmatrix} \tilde{\zeta} r_1 \\ \tilde{\zeta} r_1 \\ \tilde{\zeta} \end{bmatrix}, \Sigma\right).$$

Inequality (5) becomes

$$2W - \tilde{\zeta}^2(1 - r_1^2 - r_2^2 - 2r_1 r_2 r_{12}) > 0, \tag{9}$$

with

$$\mathbb{E}(W) = \tilde{\zeta}^2(1 - r_1^2 - r_2^2)$$
$$\mathrm{Var}(W) = \tilde{\zeta}^2(1 - r_1^2 - r_2^2 + 2r_1 r_2 r_{12}) = \sigma_W^2.$$

Once again condition $\mathrm{Var}(W) > 0$ yields

$$1 + 2r_1 r_2 r_{12} > r_1^2 + r_2^2 \tag{10}$$

and computing the probability of event (9) we get

$$P(\zeta, r_1, r_2, r_{12}) = 1 - \Phi(-1/2\sigma_W) = 1 - \Phi\left(-\frac{1}{2}|\zeta|\sqrt{1 - r_1^2 - r_2^2 + 2r_1 r_2 r_{12}}\right).$$

Hence, the probability of picking SNP 3 when it is causal is always greater than 0.5 and increasing with the absolute value of effect size $\zeta$ (see Figure 4).

Finally we calculate the probability of the BIC condition (6) in a similar fashion

$$P(\zeta, r_1, r_2, r_{12}, n) = 1 - \Phi\left(-\frac{1}{2}\left(\sigma_W + \frac{\ln(n)}{\sigma_W}\right)\right).$$

The probability is non-zero for some combinations of $r_1$, $r_2$, $r_{12}$ and $\zeta$, bounded below by 0.5 and increases in sample size $n$.

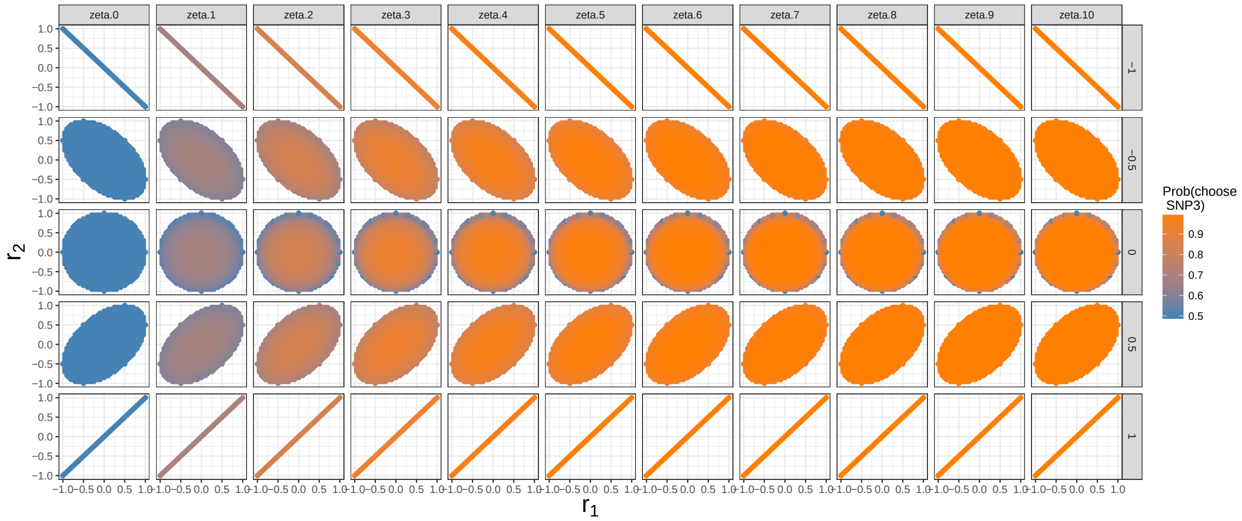

Figure 4: Probability of choosing a model under which SNP 3 is actually causal, when SNP 3 is causal, $P(\zeta, r_1, r_2, r_{12})$, under the likelihood ratio setup for varying values of $r_1$, $r_2$, different effect sizes $\zeta$ and correlation $r_{12}$ between SNPs 1 and 2 (side panels). $P(\zeta, r_1, r_2, r_{12}, n)$ is bounded below by 0.5 and is very close to 1 for all but very extreme values of $r_1$ and $r_2$ for effect sizes $\zeta > 4$.

#### 3 The ABF for a multinomial model can be expressed as a function of the ABFs of the dichotomous logistic models

We suppose we observe $N$ individuals, each individual $i$ with response $y_i \in 0, 1, \ldots, m$ for $m$ diseases and a control group, represented by 0. We assume that each individual falls into exactly one class - i.e. that no co-morbid individuals are in the sample - and that each individual has a vector of covariate data $\boldsymbol{x}_i$. A "model" is defined by which elements of $\boldsymbol{x}_i$ are used to fit a regression model to the data, which we write by replacing $\boldsymbol{x}_i$by $\boldsymbol{x}_i^M$ for model $M$. Let $\phi_{id} = \Pr(y_i = d)$ and $n_d = \sum_i I(y_i = d)$. Then the multinomial regression likelihood is defined by

$$L_M = \prod_{d=0}^{m} \prod_{i:y_i=d} \phi_{id}$$

where $\phi_{id}$ is estimated from equations

$$\log\left(\frac{\phi_{id}}{\phi_{i0}}\right) = \beta_d' \boldsymbol{x}_i^M, \quad i = 1, \ldots, N, \quad d = 1, \ldots, m$$

and $\sum_{d=0}^{m} \phi_{id} = 1$. Thus,

$$\phi_{id} = \frac{exp(\beta_d' \boldsymbol{x}_i^M)}{1 + \sum_{d=1}^{m} exp(\beta_d' \boldsymbol{x}_i^M)}$$

The corresponding logistic models have likelihoods

$$L_d = \prod_{i:y_i=d} \theta_{id} \times \prod_{i:y_i=0} (1 - \theta_{id})$$

where $\theta_{id} = \Pr(y_i = d | y_i \in \{0, d\})$ and $\log\left(\frac{\theta_{id}}{1-\theta_{id}}\right) = \gamma_d' \boldsymbol{x}_i^M$.

Begg and Gray[?] have shown that $\hat{\gamma}_d = \hat{\beta}_d$, $d \neq 0$, and that

$$\hat{\theta}_{id} = \frac{\hat{\phi}_{id}}{\hat{\phi}_{id} + \hat{\phi}_{i0}}.$$

We wish to show, comparing a specific model to the null model, that the approximate Bayes' factor (ABF) from the multinomial model is approximately proportional to the product of ABF from the logistic models. Accurate approximations to the Bayes' factor comparing the likelihood of the data under model $M$ and the null model (and integrating over the values of the regression coefficients) have been derived [?].

We use an approximation based on the Schwartz inequality which states that, writing $B_{M0}$ for the BF comparing model $M$ with the null model, and with $S$ defined as

$$S = \log \Pr(D|\hat{\theta}_M, M) - \log \Pr(D|\hat{\theta}_0, M_0) - \frac{1}{2}(k_M - k_0)\log(n),$$

$$\frac{S - \log B_{M0}}{\log B_{M0}} \to 0$$

as sample size $n \to \infty$, where $k_j$ denotes the length of the parameter vector $\theta_j$ whose maximum likelihood estimate is denoted $\hat{\theta}_j$. Thus, $S$ can be used as an ABF.

Recall we use the term "configuration" to describe a set of $m$ models, $M_1, M_2, \ldots M_m$ such that $M_k$ is the model used to describe case group $k$. Under the multinomial model, the log(ABF) can be written

$$
\begin{aligned}
B^M = & \sum_{d=1}^{m} \sum_{i:y_i=d} \hat{\beta}_d' \boldsymbol{x}_i - \sum_i \log\left(1 + \sum_{d=1}^{m} \exp(\hat{\beta}_d' \boldsymbol{x}_i)\right) \\
& - \sum_{d=1}^{m} \sum_{i:y_i=d} \hat{\beta}_{0d} + \sum_i \log\left(1 + \sum_{d=1}^{m} \exp(\hat{\beta}_{0d})\right) \\
& - \frac{1}{2}(k_C - m)\log(N)
\end{aligned}
$$

where $k_C = \sum_d k_d$ is the total number of parameters in component models $M_1, \ldots, M_m$, and $\hat{\beta}_d$ are the MLE of $\beta$ under model $M_d$ relating to disease $d$ and $\hat{\beta}_{0d}$ are the MLE of $\beta$ under the null (intercept only) model. Under the logistic model for disease $d$, the log(ABF) can be written

$$
\begin{aligned}
log(B_d^L) = & \sum_{i:y_i=d} \hat{\beta}_d' \boldsymbol{x}_i - \sum_{i:y_i \in \{0,d\}} \log(1 + e^{\hat{\beta}_d' \boldsymbol{x}_i}) \\
& - \sum_{i:y_i=d} \hat{\beta}_{0d} - \sum_{i:y_i \in \{0,d\}} \log(1 + e^{\hat{\beta}_{0d}}) \\
& - \frac{1}{2}(k_d - 1)\log(n_d + n_0)
\end{aligned}
$$

So that the difference between $B^M$ and $\sum_d B_d^L$ is

$$
\begin{aligned}
D = & \sum_i \left[\log\left(1 + \sum_{d=1}^{m} \exp(\hat{\beta}_{0d})\right) - \log\left(1 + \sum_{d=1}^{m} \exp(\hat{\beta}_d' \boldsymbol{x}_i)\right)\right] \\
& + \sum_{d=1}^{m} \sum_{i:y_i \in \{0,d\}} \left[\log\left(1 + e^{\hat{\beta}_d' \boldsymbol{x}_i}\right) - \log\left(1 + e^{\hat{\beta}_{0d}}\right)\right] \\
& + \frac{1}{2}\sum_d (k_d - 1)\log(n_d + n_0) - \frac{1}{2}\log(N)(k_C - m)
\end{aligned}
$$

Set

$$
\begin{aligned}
\eta &= \frac{1}{2}\sum_d (k_d - 1)\log(n_d + n_0) - \frac{1}{2}\log(N)\sum_d (k_d - 1) \\
&= \frac{1}{2}\sum_d (k_d - 1) log\left(\frac{n_d + n_0}{N}\right)
\end{aligned}
\tag{11}
$$

We show next that $D - \eta \simeq 0$.

Recall $\phi_{id} = Pr(y_i = d)$, and note that under model $M$, $\log \hat{\phi}_{i0} = -log\left(1 + \sum_{d=1}^{m} \exp(\hat{\beta}'_d \boldsymbol{x}_i)\right)$. Note also that $\hat{\beta}_{0d} = \log(n_d/n_0)$ so that

$$-\log\left(1 + \sum_{d=1}^{m} \exp(\hat{\beta}_{0d})\right) = \log(n_0/N), \quad \text{and}$$
$$-\log\left(1 + exp(\hat{\beta}_{0d})\right) = \log\left(n_0/(n_0+n_d)\right).$$

Thus

$$D - \eta = -N\log(n_0/N) + \sum_i \log \hat{\phi}_{i0} - \sum_{d=1}^{m} \sum_{i:y_i \in \{0,d\}} \log \frac{\hat{\phi}_{i0}}{\hat{\phi}_{i0} + \hat{\phi}_{id}} + \sum_{d=1}^{m} (n_0 + n_d) \log \frac{n_0}{n_0 + n_d}$$

Now, note that $\frac{1}{N}\sum_i \hat{\phi}_{i0} = n_0/N$. We consider $\sum_i \log(\hat{\phi}_{i0})$ as a sum of Taylor series expansions of $\log \hat{\phi}_{i0}$ about $\frac{n_0}{N}$:

$$\begin{aligned}
\sum_i \log \hat{\phi}_{i0} &\simeq \sum_i \left(\log\left(\frac{n_0}{N}\right) + \frac{N}{n_0}\right)\left(\hat{\phi}_{i0} - \frac{n_0}{N}\right) + O\left(\left(\hat{\phi}_{i0} - \frac{n_0}{N}\right)^2\right) \\
&\simeq N\log\left(\frac{n_0}{N}\right) + \frac{N}{n_0}\sum_i \left(\hat{\phi}_{i0} - \frac{n_0}{N}\right) \\
&= N\log\left(\frac{n_0}{N}\right) + \frac{N}{n_0}(n_0 - n_0) \\
&= N\log\left(\frac{n_0}{N}\right)
\end{aligned}$$

neglecting terms in $O\left(\left(\hat{\phi}_{i0} - \frac{n_0}{N}\right)^2\right)$ and smaller.

Similarly,

$$\sum_{i:y_i \in \{0,d\}} \log \frac{\hat{\phi}_{i0}}{\hat{\phi}_{i0} + \hat{\phi}_{id}} = \sum_{i:y_i \in \{0,d\}} \log \hat{\theta}_{i0} \simeq (n_0 + n_d)\log\left(\frac{n_0}{n_0 + n_d}\right)$$

Therefore,

$$D - \eta \simeq -N\log(n_0/N) + N\log(n_0/N) - \sum_{d=1}^{m}(n_0+n_d)\log\left(\frac{n_0}{n_0+n_d}\right) + \sum_{d=1}^{m}(n_0+n_d)\log\left(\frac{n_0}{n_0+n_d}\right)$$

$$= 0$$

so that

$$B^M \simeq \eta + \sum_d B_d^L \tag{12}$$

To confirm the accuracy of this approximation defined by equations (11)– (12), we simulated genetic data for varying numbers of cases (two diseases) and controls and calculated logistic ABFs for all possible models using the R package BMA. From these, we calculated the summed log ABF for all possible configurations. For comparison we calculated the log ABF for the comparitor multinomial model directly using the R package mlogitBMA. Finally, we regressed the multinomial log ABF on the summed logistic log ABFs and stored the estimated intercept slope and $R^2$ of this final linear regresion model. For each dataset, we also calculated the univariate $p$ values for each SNP and disease, and stored the minimum $p$ value for each disease. We repeated this procedure 15,000 times, and found that when the minimum $p$ value was below $10^{-7}$ that the multinomial and summed logistic log ABFs were linearly related ($R^2 \simeq 1$) with a slope of 1, indicating the the multinomial ABF could be expressed as approximately proportional to the product of the logistic ABFs (Figure 5). For larger $p$ values, the approximation was less exact, with the average slope of the regression approaching 1.02 for datasets with minimum $p$ values around 0.01, but $R^2$ remaining very high, at $> 0.9996$ for all simulations. Given that the accepted threshold for genomewide significance is typically $< 10^{-8}$, we concluded that this approximation was valid in the range of datasets for which fine mapping might be useful.

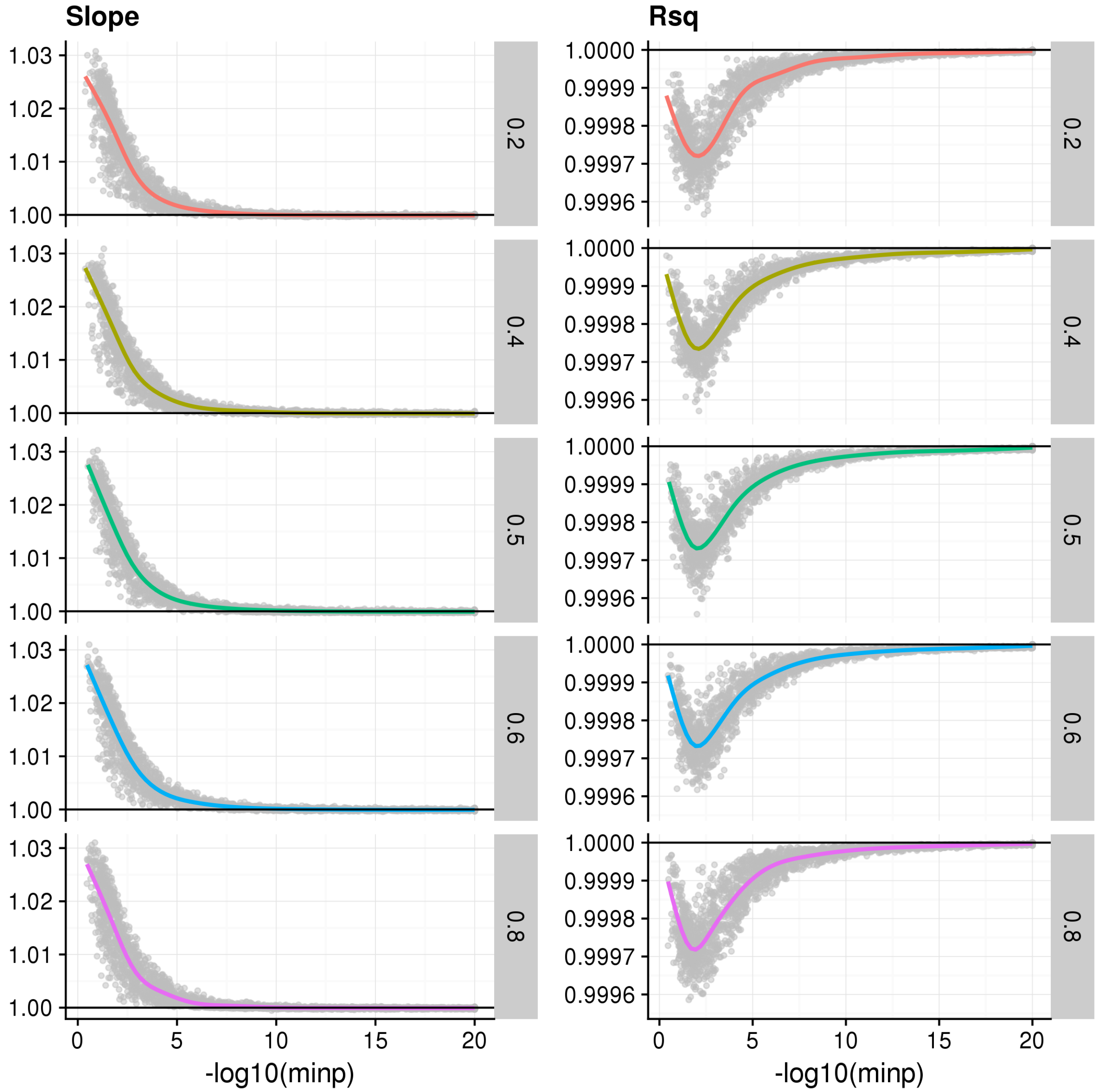


Figure 5: Comparison of log ABF for a multinomial model with the sum of log ABF for component logistic models. We simulated case-control and genetic data for varying sample sizes, effect sizes, and number of causal variants. We regressed the multinomial log ABF on the sum of logistic log ABFs and found the approximation was valid ($R^2 > 0.9996$ and slope$\simeq 1$) when the minimum p value in both datasets (x-axis) was $< 10^{-8}$. Points represent the individual estimates of slope (left column) and $R^2$ (right column). Rows are stratified according to the proportion of cases of disease 1 in the simulated sample.

### 4 Memory-efficient calculation of the marginal posteriors for each disease

Let us consider possible models $M_i$, $i = 1, \ldots, n$. Each model $i$ has a prior $p_i$, and a Bayes factor for disease 1 and 2, $b_i$ and $d_i$ respectively under a logistic model with $k_i$ parameters. We show above that the joint approximate Bayes factor for a configuration

$$C_{i,j} = M_i \text{ for disease } 1, M_j \text{ for disease } 2$$

is a function of the ABFs from the dichotomous logistic models

$$B_{ij} \propto b'_i d'_j,$$

where

$$b'_i = b_i \times \exp(\eta_1(M_i)),$$

$$d'_j = d_j \times \exp(\eta_2(M_j))$$

and

$$\eta_l(M_i) = \exp\left((k_i - 1) \times \frac{1}{2} \log\left(\frac{n_l + n_0}{N}\right)\right), \quad l = 1, 2$$

Thus, the posterior for configuration $C_{i,j}$ is

$$PP_{ij} \propto Pr(C_{i,j}) b'_i d'_j.$$

We incorporate our prior belief on shared causal variants between diseases by upweighting configurations corresponding to such sharing compared to those that don't. We set

$$Pr(C_{i,j}) = p_i p_j \kappa^{M_i \cap M_j \neq \varnothing} \tau_{ij}$$

$\kappa$ is the upweighting factor, and $\tau_{ij}$ is a normalisation factor, chosen to ensure that

$$\sum_{i:|M_i|=m,j:|M_j|=l} p_i p_j = \sum_{i:|M_i|=m,j:|M_j|=l} p_i p_j \kappa^{M_i \cap M_j \neq \varnothing} \tau_{ij} \tag{13}$$

ie, that the prior belief for a configuration corresponding to given model sizes doesn't vary with $\kappa$. The equality in (13) implies

$$\binom{n}{m}\binom{n}{l}\pi(m)\pi(l) = \tau_{ij}\pi(m)\pi(l)\left[\binom{n}{m}\binom{n-m}{l} + \kappa\binom{n}{m}\left(\binom{n}{l} - \binom{n-m}{l}\right)\right]$$

$$\tau_{ij} = \frac{\binom{n}{l}}{\binom{n-m}{l} + \kappa\left[\binom{n}{l} - \binom{n-m}{l}\right]} \tag{14}$$

for models $M_i$ and $M_j$ with sizes $m$ and $l$ respectively.

Considering the form of the marginal model posterior probabilities for each disease helps understand how $\kappa > 1$ allows information from disease 2 to be used in our inference for disease 1.

The posterior probability of $M_1$ for disease 1 is proportional to a sum of the posterior probabilities of all configurations $C_{1,j}$, $j = 1, \ldots, n$. Let $I_{i,j}$ be an indicator function, taking the value 1 if $M_i \cap M_j \neq \varnothing$ and 0 otherwise. Then

$$\begin{aligned}
Pr(M_i \text{ for disease 1}|\text{Data}) &\propto \sum_j p_i p_j b'_i d'_j \times \kappa^{I_{i,j}} \tau_{ij} \\
&= p_i b'_i \left( \sum_{j:I_{i,j}=0} \tau_{ij} p_j d'_j + \kappa \sum_{j:I_{i,j}=i} \tau_{ij} p_j d'_j \right) \\
&= p_i b'_i \left( \sum_j \tau_{ij} p_j d'_j + (\kappa - 1) \sum_{j:I_{i,j}=i} \tau_{ij} p_j d'_j \right) \\
&= p_i b'_i \left( 1 + (\kappa - 1) \frac{\sum_{j:I_{i,j}=1} \tau_{ij} p_j d'_j}{\sum_j \tau_{ij} p_j d'_j} \right)
\end{aligned}$$

Noting the similarity to

$$Pr(M_i \text{ for disease 1}|\text{Data for disease 1 only}) \propto b_i p_i$$

we can see that information from disease 2 enters by modifying the prior for model 1 according the posterior support for disease 2 for models that contain any overlap with $M_1$.

With $n > 2$ diseases, each disease may share causal variants with $n-1$ other diseases. We have more choices now, in terms of how to formulate the joint model - do we upweight further configurations that display sharing between more than 2 diseases? Given the interpretation above of the marginal posterior for one disease, we chose to focus on pairwise sharing of each of $n-1$ other diseases with a single disease of interest. This implies that, if our focus is on disease 1, we consider a prior of the form

$$\pi(C_{ijk}) \propto p_i p_j p_k \kappa^{I(M_i \cap M_j \neq \varnothing)} \kappa^{I(M_i \cap M_k \neq \varnothing)} \tau_{ij} \tau_{ik}$$

(and similar for four or more diseases). This corresponds to a marginal posterior for disease 1 (whose models are indexed by $i$)

$$Pr(M_i \text{ for disease 1}|\text{Data}) \propto p_i b'_i \left(1 + (\kappa - 1)\frac{\sum_{j:I_{i,j}=1} \tau_{ij} p_j d'_j}{\sum_j \tau_{ij} p_j d'_j}\right) \left(1 + (\kappa - 1)\frac{\sum_{k:I_{i,k}=1} \tau_{ik} p_k d'_k}{\sum_k \tau_{ik} p_k d'_k}\right)$$

This formulation also enables memory efficient calculation of the individual disease marginal posteriors, by stepping through the sums over all configurations, storing only the contents of each large bracket on the right hand side.

As before, so that the prior on any given model size is independent of $\kappa$, we have

$$\sum_{i:|M_i|=m, j:|M_j|=l, k:|M_k|=o} p_i p_j p_k = \sum_{i:|M_i|=m, j:|M_j|=l, k:|M_k|=o} p_i p_j p_k \kappa^{M_i \cap M_j \neq \varnothing} \tau_{ij} \kappa^{M_i \cap M_k \neq \varnothing} \tau_{ik}$$

which leads to

$$\begin{aligned}
\tau_{ij}\tau_{ik} &= \frac{\binom{n}{o}\binom{n}{l}}{\binom{n-m}{o}\binom{n-m}{l} + \kappa\left[\left(\binom{n}{l} - \binom{n-m}{l}\right)\binom{n-m}{o} + \left(\binom{n}{o} - \binom{n-m}{o}\right)\binom{n-m}{l}\right] + \kappa^2\left(\binom{n}{o} - \binom{n-m}{o}\right)\left(\binom{n}{l} - \binom{n-m}{l}\right)} \\
&= \frac{\binom{n}{l}\binom{n}{o}}{\left(\kappa\binom{n}{l} - \kappa\binom{n-m}{l} + \binom{n-m}{l}\right)\left(\kappa\binom{n}{o} - \kappa\binom{n-m}{o} + \binom{n-m}{o}\right)}
\end{aligned}$$

for models $M_i$, $M_j$, $M_k$ with sizes $m$, $l$, $o$ respectively, which is solved for $\tau_{ij}$, $\tau_{ik}$ as given by equation (14).

#### 5 Choice of $\kappa$

It may be hard to directly elicit values for the prior parameter $\kappa$, that upweights configurations with pairwise sharing of variants between diseases vs configurations without sharing. We set out here how a value for $\kappa$ may be derived from a quantity that may be more easily elicited - the probability that a pair of diseases share any causal variant (with either concordant or discordant direction of effect) within a region that they both show association, which we denote $P_\kappa$.

Recall the prior for a configuration specified by model $M_i$ for disease 1 and model $M_j$ for disease 2 is

$$\Pr(C_{i,j}) \propto p_i p_j \kappa^{I(M_i \cap M_j \neq \varnothing)} \tau_{ij}.$$

where

$$\tau_{ij} = \frac{\binom{n}{n_j}}{\left[\binom{n}{n_j} - \binom{n-n_i}{n_j}\right]\kappa + \binom{n-n_i}{n_j}} = \frac{\binom{n}{n_i}}{\left[\binom{n}{n_i} - \binom{n-n_j}{n_i}\right]\kappa + \binom{n-n_j}{n_i}}$$

Note that, given $n$ SNPs in a region, and using $n_i,\ n_j$ to denote the sizes of models $M_i,\ M_j$ respectively, then the number of models that can be selected with size $n_i$ is $\binom{n}{n_i}$, the number of configurations with model sizes $n_i,\ n_j$ is $\binom{n}{n_i}\binom{n}{n_j}$, and the number of these that contain no shared causal variants is $\binom{n}{n_i}[\binom{n-n_i}{n_j}$ (equivalently, the number which contain at least one shared causal variant is $\binom{n}{n_i}[\binom{n}{n_j} - \binom{n-n_i}{n_j})]$). The prior probability of no sharing in causal variant models is

$$\begin{aligned} P_0 &= \sum_m \sum_l \binom{n}{m}\binom{n-m}{l} \times \frac{\pi(m)}{\binom{n}{m}}\frac{\pi(l)}{\binom{n}{l}} \times \frac{\binom{n}{l}}{[\binom{n}{l} - \binom{n-m}{l}]\kappa + \binom{n-m}{l}} \\ &= \sum_m \sum_l \frac{\pi(m)\pi(l)}{\left[\binom{n}{l}/\binom{n-l}{m} - 1\right]\kappa + 1} \end{aligned}$$

Then, assuming our prior probability that two diseases share no causal variants in the region of interest is $F_0$, $\kappa$ may be found by numerically solving the equation

$$\frac{F_0}{1-F_0} = \frac{P_0}{1-P_0}$$

which can be set to any elicited value, and numerically solved for $\kappa$.

For $d > 2$ diseases, $P_0$ becomes

$$P_0(d) = \sum_m \pi(m) \left( \sum_l \binom{n-m}{l} \frac{\pi(l)}{\left[\binom{n}{l} - \binom{n-m}{l}\right] \kappa + \binom{n-m}{l}} \right)^{d-1}$$

but we have to be careful about specifying the prior probability for no pairwise sharing. If the other diseases were totally independent, a natural prior value would be $F_0^{d-1}$. If the other diseases were totally dependent, then the prior would remain at $F_0$. In the absence of strong prior knowledge about this, we suggest that $F_0^{\sqrt{d-1}}$ is a sensible compromise, but that both extreme values $F_0$ and $F_0^{d-1}$ should also be explored.
