## Supplementary Figures for "Sharing information between related diseases using Bayesian joint fine mapping increases accuracy and identifies novel associations in six immune mediated diseases"

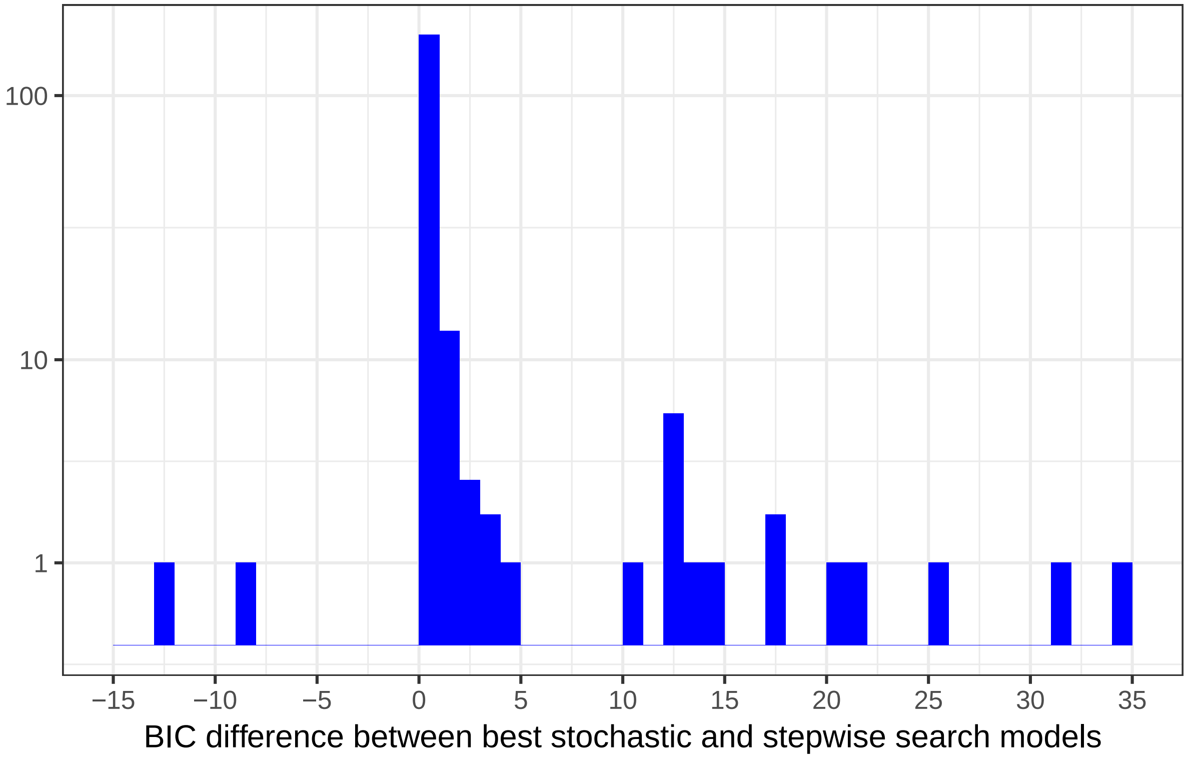


Supplementary Figure 1: Histogram of BIC(Stepwise Search) - BIC(Stochastic Search) for the best stochastic and stepwise search models from each region-disease analysis. Stochastic search tends to have a BIC that is the same or smaller as that from stepwise search, indicating a fit that is at least as good or better than stepwise search.The BIC is calculated using the best SNP model from stochastic search, and for two regions, the null model is preferred over each individual SNP model, coinciding with the two instances of stepwise appearing to select a better-fitting model than stochastic search. However, in these two instances, the best non-null model selected by stochastic search agrees with that of stepwise search.

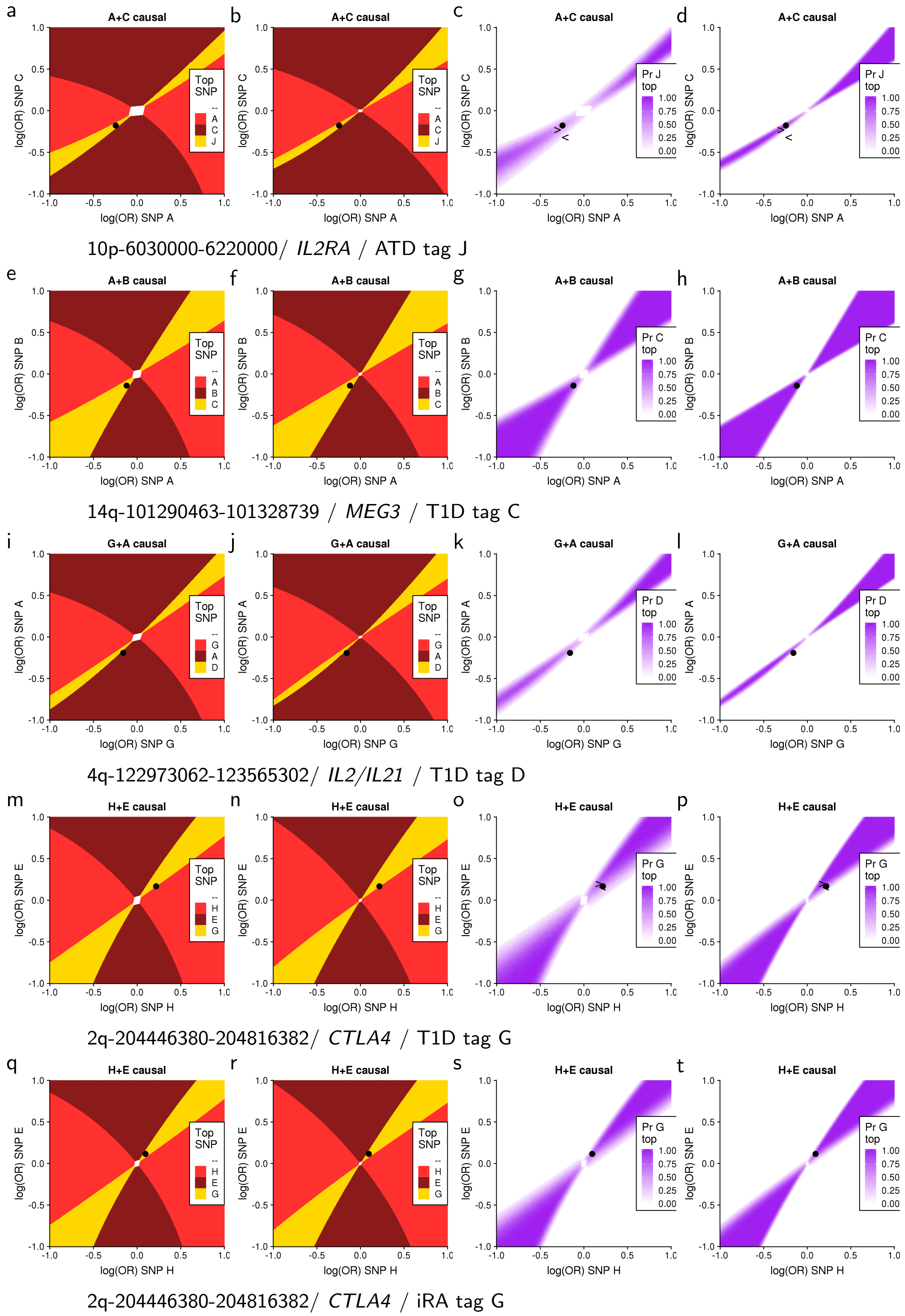


Supplementary Figure 2: Each plot **a-t** is a 2x2 grid, where each point represents a single two SNP causal model, specified by effects (log odds ratio) for the causal SNPs on the X and Y axes. First two columns: "Sunbeam plots" show which SNP is expected to be selected first in a step wise search at each position. Last two columns: "Probability plots" show the probability that the tag SNP was first selected in a stepwise search. Columns 1, 3: Sample sizes from relevant disease dataset; Column 2,4: sample size of 50000 cases and 50000 controls. Two sample sizes are shown so the dependence on small sample size can be evaluated. The black dot shows where the observed disease data lie.

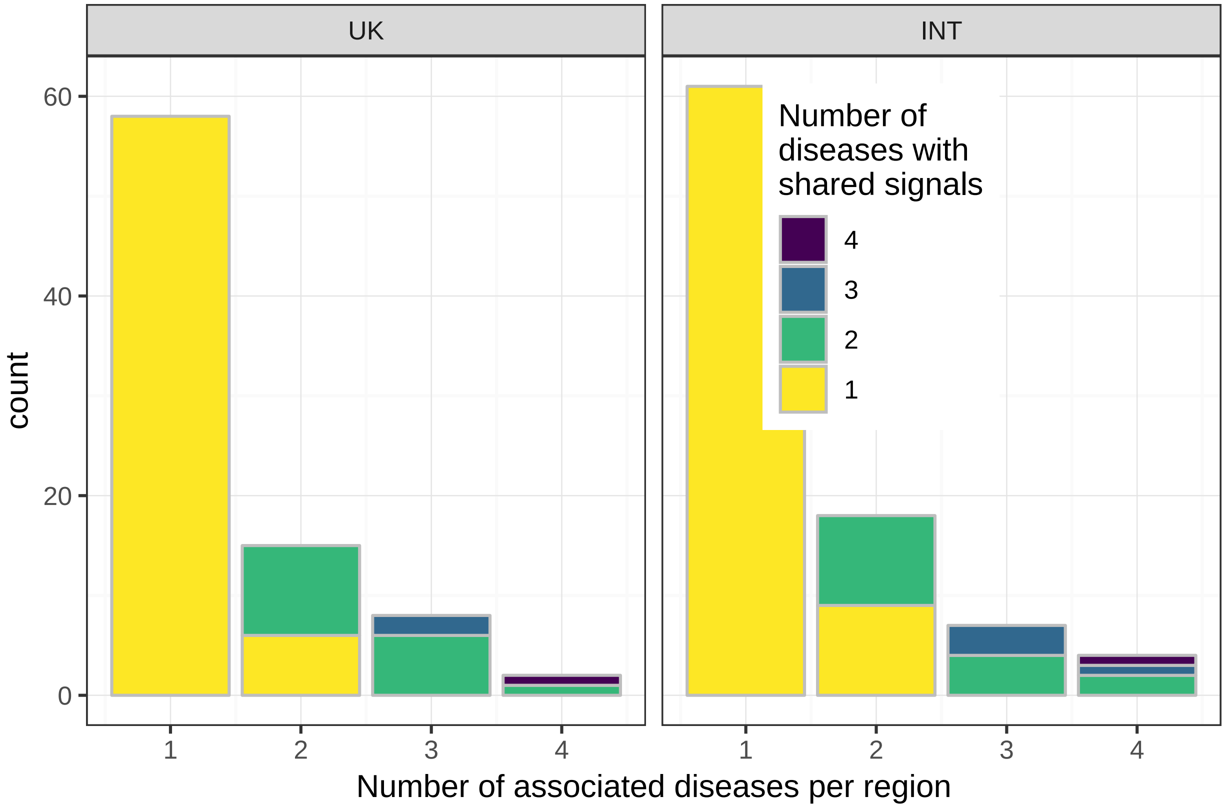


Supplementary Figure 3: Frequency of he number of associated diseases per region, partitioned by whether signals are shared between diseases, for UK samples (left) and international samples (right). We consider a signal to be shared when there exists a SNP group with MPPI ¿ 0.5 for more than one disease.
